# Simulation Framework for Generating Intratumor Heterogeneity Patterns in a Cancer Cell Population

**DOI:** 10.1101/109801

**Authors:** Watal M. Iwasaki, Hideki Innan

## Abstract

As cancer cell populations evolve, they accumulate a number of somatic mutations, resulting in heterogeneous subclones in the final tumor. Understanding the mechanisms that produce intratumor heterogeneity (ITH) is important for selecting the best treatment. Although some studies have involved ITH simulations, their model settings differed substantially. Thus, only limited conditions were explored in each. Herein, we developed a general framework for simulating ITH patterns and a simulator (tumopp). Tumopp offers many setting options so that simulations can be carried out under various settings. Setting options include how the cell division rate is determined, how daughter cells are placed, and how driver mutations are treated. Furthermore, to account for the cell cycle, we introduced a gamma function for the waiting time involved in cell division. Tumopp also allows simulations in a hexagonal lattice, in addition to a regular lattice that has been used in previous simulation studies. A hexagonal lattice produces a more biologically reasonable space than a regular lattice. Using tumopp, we investigated how model settings affect the growth curve and ITH pattern. It was found that, even under neutrality (with no driver mutations), tumopp produced dramatically variable patterns of ITH and tumor morphology, from tumors in which cells with different genetic background are well intermixed to irregular shapes of tumors with a cluster of closely related cells. This result suggests a caveat in analyzing ITH data with simulations with limited settings, and tumopp will be useful to explore ITH patterns in various conditions.

**Author Summary:** Understanding the mechanisms that produce intratumor heterogeneity (ITH) is important for selecting the best treatment. Despite a growing body of data and tools for analyzing ITH, the spatial structure and its evolution are poorly understood because of the lack of well established theoretical framework. Herein, we provide a general framework for simulating ITH patterns, under which a simulator (tumopp) is developed. Tumopp offers many setting options so that simulations can be carried out under various settings. Simulations using tumopp demonstrate that dramatically variable patterns of ITH and tumor morphology can be produced depending on the model setting. The present work provides a guideline for future simulation studies of cancer cell populations.

## Introduction

Tumors begin from single cells that rapidly grow and divide into multiple cell lineages by accumulating various mutations. The resulting tumor consists of heterogeneous subclones rather than a single type of homogeneous clonal cells [1–4]. This phenomenon is known as intratumor heterogeneity (ITH) and is a significant obstacle to cancer screening and treatment. Thus, understanding how tumors proliferate and accumulate mutations is essential for early detection and treatment decisions [5–8]. Multiregional and single-cell sequencing are promising way for uncovering the nature of ITHs within tumors [9–11], and a large amount of high-throughput sequencing data have been accumulating [12, 13] together with bioinformatic tools to interpret such data [14, 15]. However, the spatial structure and its evolution are still poorly understood [16] because of the lack of well established theoretical framework. Although some studies have involved ITH simulations, their model settings differed substantially [9, 17–21]. The purpose of the current study was to develop a general framework for simulating ITH patterns in a cancer cell population to explore all possible spatial patterns that could arise and under what conditions. To do so, we aimed to ensure that simulations do not take a very long time so that it can be used within the framework of simulation-based inference as outlined in Marjoram et al. [22] (see also refs therein).

Of the various types of cancer cell growth models, single-cell-based models are more appropriate for our purposes than continuum models that treat tumors as diffusing fluids. There are two major classes of single-cell-based models, on- and off-lattice. The former assumes that each cell is placed in a space with discrete coordinates, while the latter defines cells in more complicated ways. The current study highlights on-lattice models because they do not involve as large amounts of computation as off-lattice models. Even in simple settings, off-lattice models represent cells as spheres in a continuous space, whose position is affected by attractive and repulsive interactions with other cells [23]. Other examples include immersed boundary model [24] and subcellular element model [25], which define cells by modeling a plasma membrane and network of particles, respectively. On-lattice models define cells as either single or multiple nodes on a lattice. The cellular Potts model [26–28] is a multiple node-based on-lattice model in which a cell is represented by several consecutive nodes. This model is similar to the subcellular element model in that complicated cell shapes can be defined. In contrast, single node-based on-lattice models assume that a cell is represented by a single node on the lattice and, thus, can be considered as a kind of cellular automaton model. The computational load can be minimized with this one-by-one relationship between cells and nodes.

Of the several cellular automaton models available for cancer cell growth [9, 17–21], most are quite simple and can be readily used for simulation-based inference of parameters in cancer cell growth. These models generally consider simple patterns of cell behavior; cells can produce new cells (cell division), die or migrate somewhere else, and each cell’s behavior can be stochastically determined depending on its own state and that of its neighbors. However, there are substantial differences in model settings among previous studies, and how these differences affect the final outcome is poorly understood. Herein, we developed a general framework for simulating cellular automaton models of tumor growth called tumopp. We made our framework as flexible and reasonable as possible for on-lattice models in which each cell is located on a single node, and normal cells and extracellular matrix surrounding the tumor cells are ignored. Moreover, the environment is independent of the configuration and dynamics of the tumor cells. In other words, while tumor growth does not change the surrounding environment, its growth is affected by the environment. These conditions are commonly assumed in most previous studies [9, 17–21].

Even with these conditions for minimizing computational load, our framework is flexible enough to incorporate various factors that determine the rates of cell birth and death and how a new daughter cell is placed in the lattice. Therefore, most previous models can be described within our framework. Using our framework, we explored the effect of model settings on various aspects of the final tumor. Because some settings can have rather large effects, particularly on the spatial distribution of heterogeneous cells (i.e., ITH), it is important to choose a model that best suits the specific properties of the focal cancer being investigated. Overall, the present work provides a guideline for future simulation studies of cancer cell populations.

## Model

### General Framework of tumopp

Tumopp was developed to enable fast simulation of tumor growth by assuming (i) a cell occupies a single node in the lattice, (ii) normal (noncancer) cells are not simulated, (iii) extracellular matrix surrounding the tumor is ignored, and (iv) the environment is not affected by changes in the configuration of the tumor. The initial state could be either one or multiple tumor cells distributed in a two-dimensional (2D) or 3D lattice. The entire process can be handled step by step. Suppose there are *N*_*t*_ number of tumor cells at time *t*, and *E*_global;*t*_ denotes the global environment at time *t*. The system waits for the next event (birth, death, or migration) of one of the *N*_*t*_ cells or any kind of environmental change. Potential events that cause environmental changes include medical treatments and angiogenesis. The time to the next environmental change, *w*_*E*_, can be determined either randomly or arbitrarily. The waiting times for birth (*w*_*b,i*_), death (*w*_*d,i*_), and migration (*w*_*m, i*_) events for the *i*th cell are random variables that depend on the status of each cell.

The system proceeds from time *t* by an increment of ∆*t*. If *w*_*E*_ is smaller than any other waiting time, then ∆*t* = *w*_*E*_ is given, and the environmental change is implemented at time *t* + ∆*t*. Then, *w*_*b, i*_, *w*_*d, i*_, and *w*_*m, i*_ will all be re-evaluated under the new environment. Otherwise, no environmental change occurs during Δ*t* = min(*w_b_*,_1_,…, *w_b,Nt_*, *w_d_*,_1_,…, *w_d,Nt_*, *w_m_*,_1_,…, *w_m,Nt_*), so that the next event is cell division, death, or migration (Fig. 1). If *w*_*b,i*_ is the smallest, the next event is division of the *i*th cell. While one of the two daughter cells stays as it is, the other is placed at an adjacent node. The cell division event might involve genetic changes or differentiation of the daughter cells that could result in an increase or decrease in the ability of cell division. In the *N*_*t*_ = 3 example shown in Fig. 1A, because the minimum waiting time is *w*_*b*,2_ (in blue), the second cell undergoes cell division. In a case where *w*_*d, i*_ is the smallest, the next event is the death of the *i*th cell, and the cell is removed from the lattice. If *w*_*m, i*_ is the smallest, the next event is migration of the *i*th cell. The *i*th cell may simply move to an empty neighbor site or result in a position swap with an adjacent cell. Thus, this procedure allows simulation of a tumor growth pattern once *w*_*b, i*_, *w*_*d, i*_, and *w*_*m, i*_ are determined for all cells (see Fig. 1 for details).

**Fig 1.**
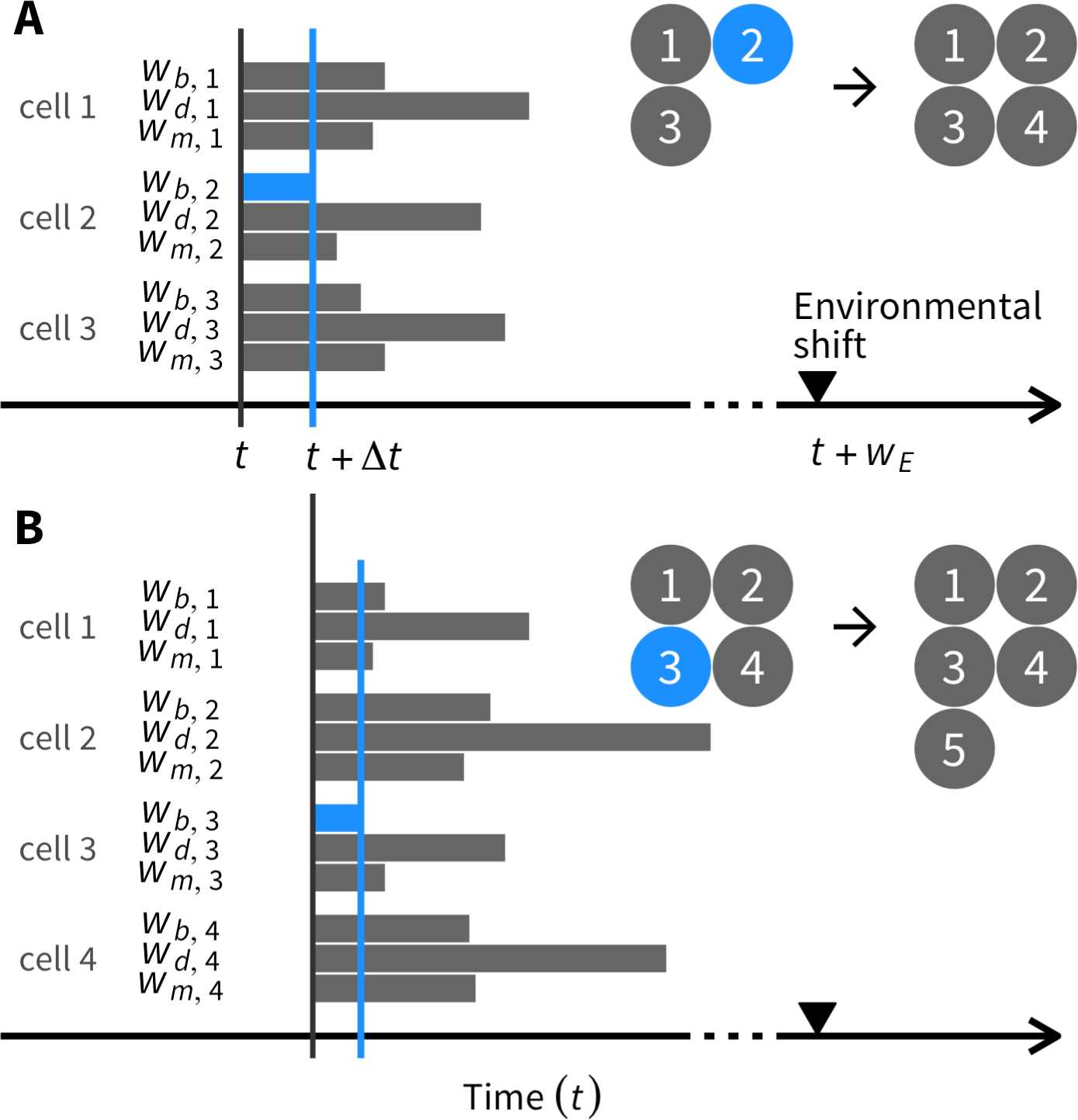
Illustration of the simulation algorithm for determining the next event. (A) An example with three cells, 1, 2, and 3 (*N_t_* = 3). The three waiting times are randomly generated for each cell as elaborated in the main text. Because *w_b_*_,2_ is the smallest (blue), the next event is cell division of the second cell, which gives birth to the fourth cell. (B) Again, the waiting times are computed for all four cells. Note that the waiting times have to be newly generated for second and fourth cells that just experienced a cell division, whereas we can reuse the waiting times for the first and third cells with ∆*t* subtracted. Because *w*_*b*,3_ is the smallest (blue), the next event is cell division of the third cell, creating the fifth cell.

*w*_*b, i*_, *w*_*d, i*_, and *w*_*m, i*_ may be random variables from certain probability density functions (PDFs), which should be flexible enough to incorporate a number of factors. These PDFs should reflect both internal cell status (*C*_*i*,*t*_) and external environment (*E*_*i*,*t*_) for the *i*th cell at time *t*. *C*_*i*,*t*_ includes various genetic and nongenetic factors:

**C1** Cell types with different proliferation potential (e.g., cancer stem cells [CSCs], transient amplifying cells [TACs], or terminally differentiated cells [TDCs]).

**C2** Genetic basis of malignancy, including the potential of cell division and death (e.g., driver mutations that have accumulated in the cell). This should also be related to the rate of migration (invasion) into nearby tissues.

*E*_*i, t*_ represents environmental factors that may be classified into two categories:

**E1** The global environment that affects the entire tumor.

**E2** The local environment within the tumor, mainly due to surrounding cancer cells.

*E*_*i, t*_ should be determined by the joint effects of various factors including E1 and E2, which may not be mutually exclusive to one another. In addition to *C*_*i, t*_ and *E*_*i, t*_, the cell status in the cell cycle may play an important role (see below for cell cycle treatment).

### Modeling with simplifying assumptions

The above framework is designed to be flexible enough to incorporate various factors, but making the model too complex would involve a substantial amount of simulation time. Here we provide several assumptions to simplify the process while keeping the model in tumopp as biologically reasonable as possible. First, we defined the simulation space, which is either regular (square) or hexagonal in 2D or 3D space (Fig. 2). The neighborhood, or adjacent sites, must also be defined because it is involved in the algorithms that determine how new cells are placed. In a regular lattice (Fig. 2), there are at least two methods to define the neighborhood. The Moore neighborhood assumes that each cell has 8 and 28 neighbors in 2D and 3D lattices, respectively, whereas the von Neumann neighborhood assumes only 4 and 6 neighbors, respectively. In the current work, we use the Moore neighborhood as in previous studies, unless otherwise mentioned. The von Neumann neighborhood assumes unrealistic behavior, thereby creating a strange tumor shape (see Discussion). The situation is simpler in a hexagonal lattice, where each cell has 6 and 12 neighbors in 2D and 3D lattices, respectively. It should be noted that there are two versions of a 3D hexagonal lattice, hexagonal close-packed and face-centered cubic. Because the difference is very small, we used the latter in the present study, which is computationally a little more tractable.

**Fig 2.**
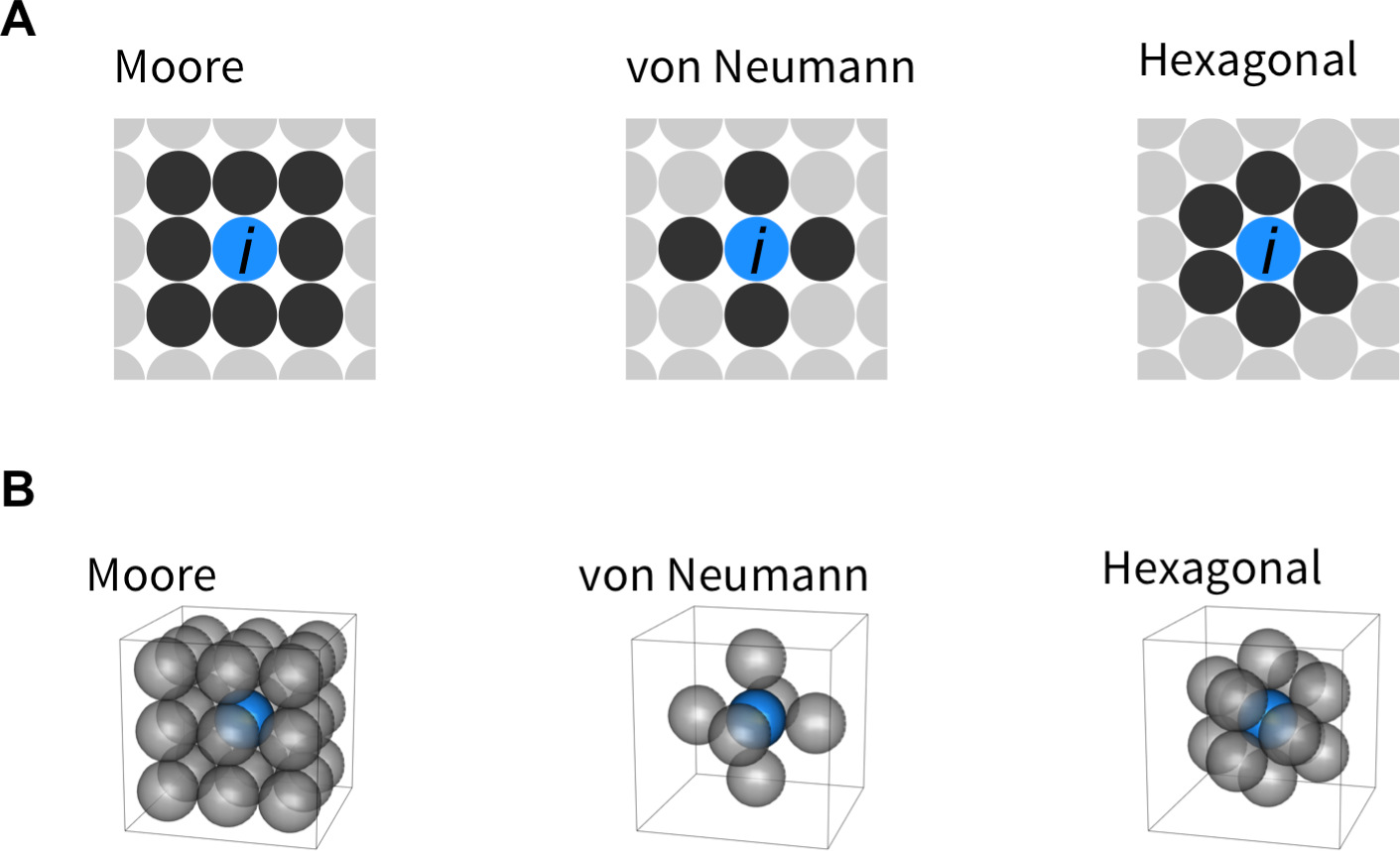
Definitions of neighborhood, or adjacent sites, in 2D (A) and 3D space (B). The focal site (*i*th cell) is shown in blue, and its adjacent sites are in black. Note that there are not multiple definitions of neighborhood in a hexagonal lattice.

The simulation process consists of a large number of steps, at which one of the cells undergoes birth, death, or migration in the simulation space. As described above (Fig. 1), the event is determined by generating random variables for waiting times (*w*_*b, i*_, *w*_*d, i*_, and *w*_*m, i*_) from certain PDFs. In this section, we describe how to model the process and determine these PDFs denoted by *f_b,i_*(*w_b,i_* | *C_i,t_, E_i,t_*), *f_d,i_*(*w_d,i_* | *C_i,t_*, *E_i,t_*), and *f_m,i_*(*w_m,i_* | *C_i,t_*, *E_i,t_*).

### Modeling waiting times

A gamma function is useful for handling the three waiting times (*w*_*b, i*_, *w*_*d, i*_, and *w*_*m, i*_) for the *i*th cell. First, consider the waiting time for cell division (*w*_*b, i*_). Suppose that the *i*th cell is a newborn cell that has just undergone cell division at time *t*. We assume that the time to the next environmental shift (*w*_*E*_) is very long (i.e., the environment is constant on the cell division time scale). Thus, the waiting time for the next cell division can be assumed to follow a gamma function:

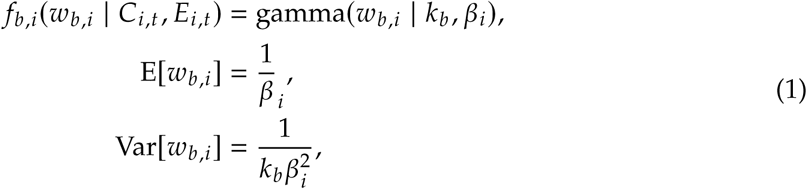

Where *f_b,i_*(*w_b,i_* | *C_i,t_*, *E_i,t_*) can be specified by only two parameters: (1) birth rate (β_*i*_), which is the reciprocal of the mean waiting time of cell division since the last cell division and referred to as the *potential* birth rate because it applies only to a newborn cell (see below for details); and (2) the shape of the distribution (*k*_*b*_). If *k*_*b*_ = ∞ is assumed, Equation 1 is given by a delta function 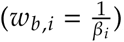
as *k*_*b*_ decreases, the distribution spreads around the mean, 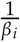
 and is identical to an exponential distribution with parameter 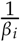
 when *k*_*b*_ 1 (Fig. 3).

A relatively large *k*_*b*_ might provide a reasonable PDF considering the cell cycle illustrated in Fig. 4. A cell has to go through interphase to get to metaphase, during which cell division occurs. This is why Equation 1 can only be applied to a newborn cell. For a cell that experienced the last cell division *t* = τ before, Equation 1 should be modified as follows:

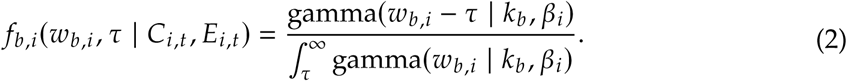

It should be noted that most previous studies [9, 17–21] ignored this effect of the cell cycle and used an exponential distribution (*k*_*b*_ = 1) instead, where extremely short cell division after the previous one is allowed. As demonstrated in Results, this simplification has a non-negligible effect on many features in simulated tumors.

Similarly, the waiting times for death (*w*_*d, i*_) and migration (*w*_*m, i*_) of the *i*th cell may be described with gamma distributions:

**Fig 3.**
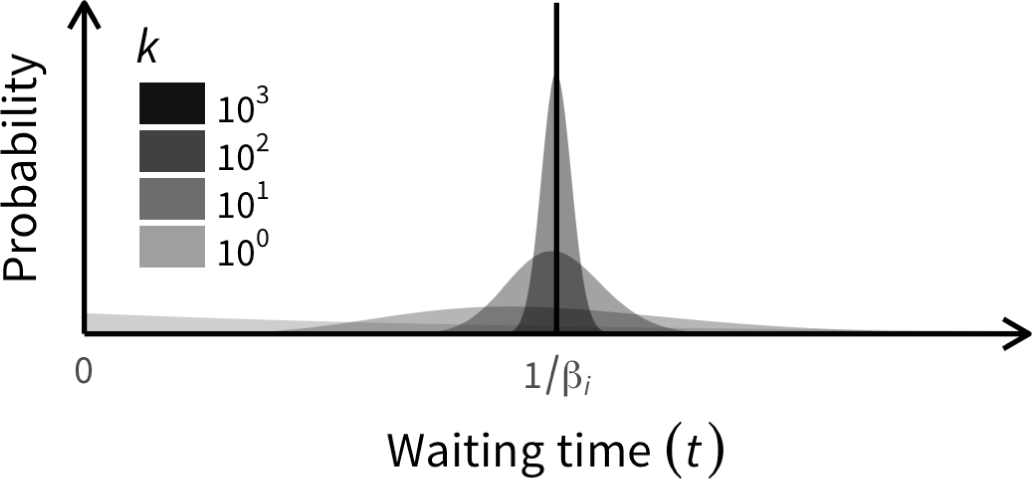
Effect of shape parameter (*k*) on gamma distribution with mean. 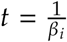. When *k* is very large, the variance of *t* is very small; when *k* is small, *t* has a wide distribution. In the extreme condition where *k* = 1, the distribution is identical to the exponential distribution with mean 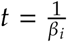.

**Fig 4.**
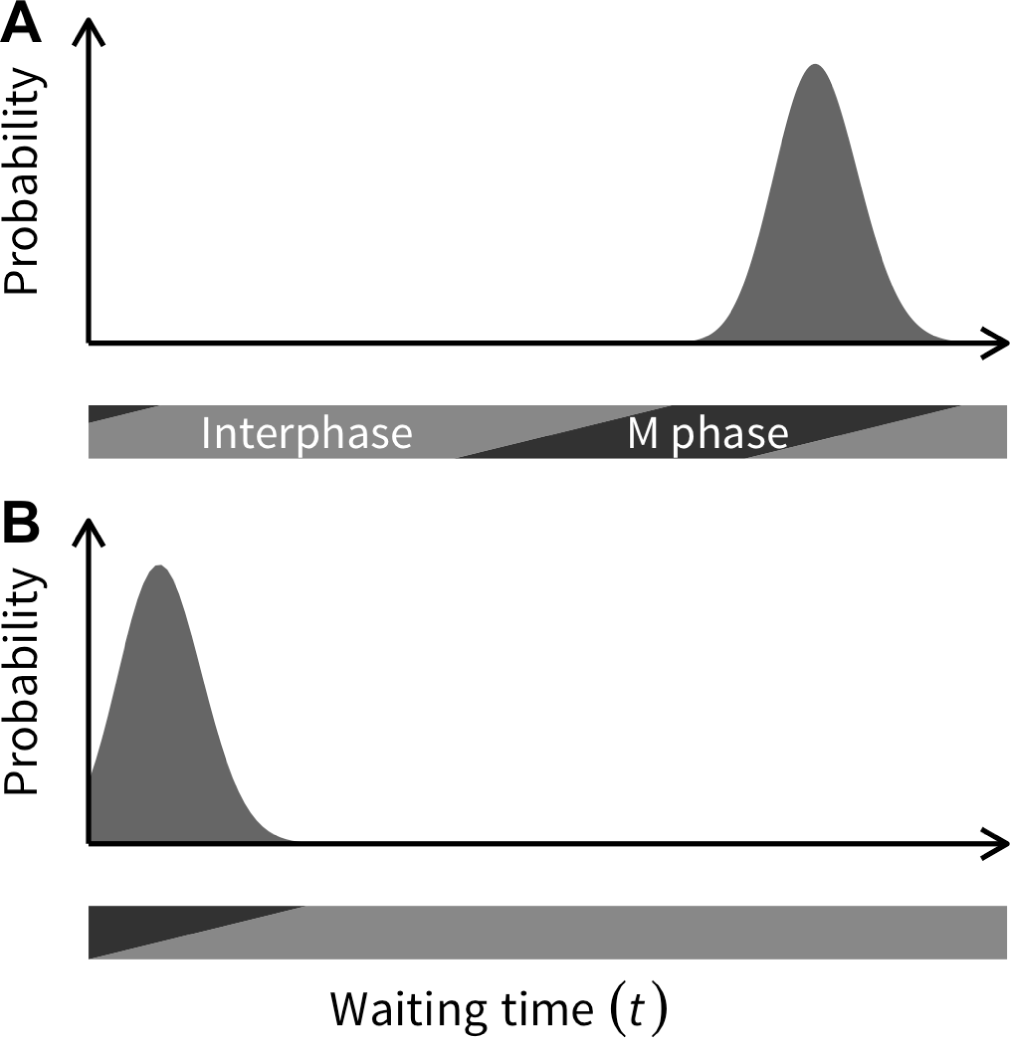
Illustrating the biological background behind using a gamma distribution with a reasonably large *k*. When a cell undergoes division, its daughter cells should enter interphase, during which they prepare for the next cell division, and it should be difficult to predict a cell division in early interphase (see text for details).

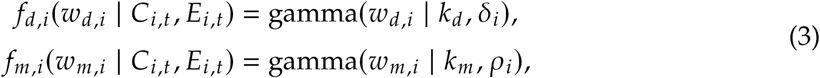

Where δ_*i*_ and ρ_*i*_ are the expected *w*_*d, i*_ and *w*_*m, i*_, respectively. In contrast to cell division, cell death and migration may not have a clear correlation with the cell cycle. If so, an exponential distribution may be used (by assuming *k*_*d*_ = 1 and *k*_*m*_ = 1 in the equations above). An exponential distribution does not require adjustment in the cell cycle (i.e., τ) because the following equation holds, which reduces the computational load:

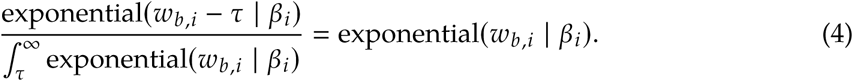

There is also an alternative treatment for cell division and death [17, 19]. Cell death might occur when the cell gets into metaphase and tries to undergo cell division but fails [29]. This can be modeled such that *w*_*b, i*_ and *w*_*d, i*_ follow a single PDF (i.e., a gamma distribution), and the outcome could be randomly assigned to cell division and death with probabilities 1 − α_*i*_ and α_*i*_, respectively. Tumopp implements these two alternative treatments. Thus, the PDFs for the three waiting times can be given once the potential rates (β_*i*_, δ_*i*_, and ρ_*i*_) are determined (see below).

### Potential birth rate

β_*i*_ should be determined by genetic and environmental factors. To incorporate the effects of the two genetic (C1 and C2) and two environmental (E1 and E2) factors, we define β_*i*_ as:

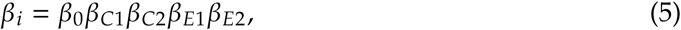

Where β_0_ is a constant value shared by all cells. β_*C*1_, β_*C*2_, β_*E*1_, and β_*E*2_ are the coefficients determined by the above-mentioned factors that constitute C1, C2, E1, and E2, respectively.

**C1** The proliferation potential of a cell largely depends on the cell types, including CSCs, TACs, and TDCs. This can be implemented through subsequent asymmetric cell divisions [19, 30]. In a simple setting, CSC can be assumed to produce another CSC with probability *p*_*s*_, and divides asymmetrically to produce a TAC with probability 1 − *p*_*s*_. A TAC has limited proliferation capacity. With *ω* as the number of cell divisions allowed for a TAC and *ω*_max_ as the maximum number of cell divisions for an initial TAC, an initial TAC has *ω*_max_, and *ω* decreases by one when it undergoes cell division. Then, the TAC becomes a TDC when *ω* reaches zero. Under this setting, it may be reasonable to assume β_*C*1_ = 1 for a CSC and TAC with *ω* > 0, and *β*_*C*1_ 0 for a TDC. Previous models with a single-cell type with unlimited proliferation potential [17, 18] can be considered a special case with *p*_*s*_ = 1 for all cells.

**C2** The rate of cell division should be largely affected by driver mutations, which may be incorporated as follows. Driver mutations are assumed to occur at rate μ per cell division. Suppose the *i*th cell has accumulated *M* driver mutations. Here, we define a driver mutation such that it affects the birth, death, and/or migration rates, either positively or negatively, and the relative effects on the three rates are denoted by *s*_β_, *s*_δ_, and *s*_ρ_ (*s*_δ_ and *s*_ρ_ are relevant to death and migration rates as explained below). Then, assuming the effects of driver mutations are additive, β_*C*2_ may be written as follows:

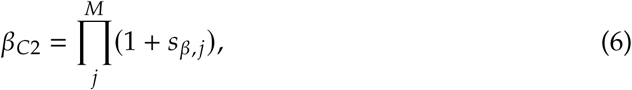

where *s*_β,*j*_ is the relative effect of the *j*th driver mutation. *s*_β, •_ may be given by a random variable from a certain distribution. Herein, we use a Gaussian distribution 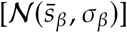
 where 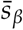
 and σ_β_ are the mean and standard deviation of the distribution, respectively.

**E1** The behavior of cancer cells should depend on their surrounding environment. For example, cells close to a nutrient source may have higher cell division rates. This might apply to cells that are close to the outer layer of the tumor or blood vessels. If so, the proliferation potential may be given by a decreasing function of the distance from these surfaces and/or blood vessels. In contrast, cell divisions will be suppressed when an anticancer drug is given. Thus, the birth rate of a cell may be given by a function of its position in the lattice:

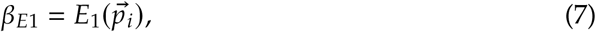

Where 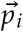
 is the position [i.e., 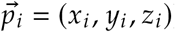
 if we set a 3D lattice. Here, we assume E1 accounts for the environment without considering the interaction between nearby cells, and the local resource competition among nearby cells is included in E2 (see below). For simplicity, tumopp assumes a uniform environment across the whole tumor. The environment might change over time, especially when a medical treatment is introduced. In our model, such an environmental change is incorporated arbitrarily, and the effect of an environmental change on each cell might depend on its genotype (i.e., configuration of driver mutations).

**E2** Growing cells are in resource competition because cell proliferation should depend on resources, such as space, oxygen, and other nutrients. It should be noted that this factor is not mutually exclusive with E1. Because competition may correlate with local density, β_*E*2_ can be given by where ϕ_*i*_ is the proportion of empty nodes in the adjacent sites of the *i*th cell. In practice, tumopp employs three models to incorporate this factor:

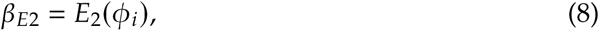

**Constant-rate model** where the birth rate is constant regardless of the availability of empty sites (ϕ_*i*_).

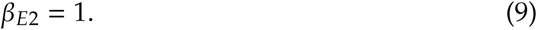

**Step-function model** where birth rate is given by a Heaviside function of ϕ_*i*_ such that cell division can occur only when there is at least one empty site available around the *i*th cell.

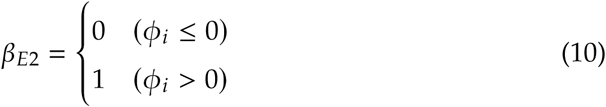

**Linear-function model** where birth rate is proportional to the number of empty neighbors [18].

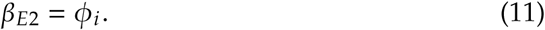

### Death rate

Similar to the birth rate case, we can define the potential death rate as:

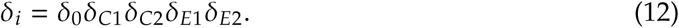

The situation may not be as complicated for the death rate as with the birth rate. C1 and E2 may not be very relevant if we consider that cell death occurs simply by chance regardless of cell type or local environment (δ_*C*1_ = δ_*E*2_ = 1 is assumed in tumopp). C2 should play a crucial role because some driver mutations significantly reduce the death rate (e.g., by avoiding apoptosis). By assuming all mutation effects are additive, this effect can be incorporated using Equation 6 with *s*_β_ replaced by *s*_δ_. Environmental changes (E1) are incorporated arbitrarily following the birth rate.

### Migration rate

The potential migration rate is given by Similar to the death rate, C2 should be most relevant to the migration rate because some mutations may provide higher mobility to the host cell (e.g., by changing adhesion molecules on membranes). Again, Equation 6 can be used here with *s*_β_ replaced by *s*_ρ_. C1 and E2 are ignored, and E1 is incorporated arbitrarily (see above).

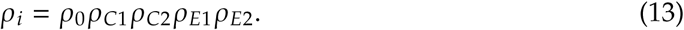

## Treatment of cell division, death, and migration in a lattice

Cell division produces two daughter cells. When placing these two cells in a lattice, we assume that one of them stays at the original site. There are several methods for placing the other cell. Tumopp employs four push methods following previous studies, which are explained by assuming that cell division occurs at (*x*, *y*, *z*) in a 3D lattice. We first describe the four methods assuming the constant-rate model, followed by their behavior in the step-and linear-function models.

**Push method 1** One new daughter cell is placed randomly on one of the adjacent neighboring sites (Fig. 2 defines adjacent sites). The direction to the adjacent site in which the cell is placed is randomly determined; for example, if the direction increases the value of *x*, then the daughter cell is placed at (*x* + 1, *y*, *z*). If (*x* + 1, *y*; *z*) is already occupied, the pre-existing cell is moved in the same direction to (*x* + 2, *y*, *z*). If a cell has already occupied (*x* + 2, *y*, *z*), then it is further shifted to (*x* + 3, *y*, *z*). Thus, the succeeding movement is repeated along in the same direction until no more push is needed. This model is used by Sottoriva et al. [17].

Push methods 2–4 are different from 1 in that if there are any empty adjacent neighboring sites available, a new daughter cell is placed to fill one of them. When no empty sites are available, methods 2–4 differ in the way they determine which neighboring cell to push out. All of the push methods use statistic *l*_min_, the minimum distance (the number of consecutive occupied sites) to the nearest empty site for a specific direction. If we assume the Moore neighborhood (Fig. 2), it is computed in all of 26 possible directions.

**Push method 2** The push direction is randomly determined, and the probability for each direction is weighted by 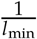
 Once the direction is determined such that the direction increases the value of *x*, for example, the daughter cell is placed at (*x* + 1, *y*, *z*). If (*x* + 1, *y*, *z*) is already occupied, the pre-existing cell is moved in the same direction to (*x* + 2, *y*, *z*). If a cell has already occupied (*x* + 2, *y*, *z*), then it is further shifted to (*x* + 3, *y*, *z*). Thus, the succeeding movement is repeated in the same direction, such that *l*_min_ cells are automatically pushed toward the surface. This method was adopted by Uchi et al. [9].

**Push method 3** The new cell is placed at the adjacent site in the direction with the smallest *l*_min_. At that site, *l*_min_ for the pre-existing cell is again computed in all directions, and the pre-existing cell is moved one step in the direction with the smallest *l*_min_. This process is continued until an empty site is found so that no more push is needed. This method is according to model C of Waclaw et al. [18].

**Push method 4** Simplified version of push methods 2 and 3, wherein the push direction is determined only once with the smallest *l*_min_. Then, *l*_min_ cells in a row are all pushed toward the surface as described for push method 2.

Thus, tumopp implements four push methods in combination with the constant-rate model, whereas the situation is much simpler in the step- or linear-function models that assume only cells with empty sites available in the neighborhood can undergo cell division. Thus, there are virtually only two distinct methods; push method 1 also works in the step- or linear-function models, while the behavior of push methods 2–4 are identical. This is because one of the empty sites in the neighborhood is automatically filled by a new cell, otherwise no cell division occurs (with no empty sites available), and how a pre-existing cell is pushed is irrelevant.

For cell death, the cell simply disappears, and the node becomes empty, while migration is defined as a single-step move of a cell in the lattice. Suppose that the cell at (*x*, *y*, *z*) is migrant and moves to (*x*, *y*, *z* + 1). If (*x*, *y*, *z* + 1) is empty, the cell simply moves and (*x*, *y*, *z*) becomes empty. If there is a pre-existing cell at (*x*, *y*, *z* + 1), the cells at (*x*, *y*, *z*) and (*x*, *y*, *z* + 1) are replaced by each other.

## Simulation

Tumopp was developed as a simulator for generating patterns of cancer cell growth under the setting described in the previous section. Table 1 summarizes the options and parameters involved in tumopp. It is first necessary to set either a regular (square) or hexagonal lattice in 2D or 3D space. Then, an initial cell is placed at position 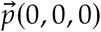
 in 3D space or 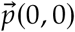
 in 2D space. The initial cell has to be a stem cell (CSC) with *ω* = *ω*_max_. This initial cell and its descendants undergo cell division, death, and migration. Their rates are determined by Equations 5, 12, and 13, respectively; all parameters involved are summarized in Table 1.

Our model is flexible so that most previous models can be described in our framework; Table 1 compares our model with four representative simulation studies on ITH. For example, while all previous studies used a regular lattice for the simulation space, we added a hexagonal lattice. We believe a hexagonal lattice is biologically more reasonable because the distance to all neighbor cells is identical. Following Poleszczuk et al. [19], our model has a flexible setting for different cell types, from CSC to TDC with intermediate states, although the other three studies assumed that all cells are CSCs (i.e., *p*_*s*_ = 1 is fixed). In our model, the rates of cell division, death, and migration are defined such that a number of factors are taken into account, while the four previous studies only incorporated part of them. Moreover, our model includes all of the methods for placing a new daughter cell that were used in the four previous studies. Tumopp is unique because it employs a gamma function for *w*_*b, i*_, while all four previous studies used an exponential or geometric function. Both are essentially identical, simple decreasing functions, except that an exponential function is continuous while a geometric function is discrete. Note that an exponential function is a special case of a gamma function with the shape parameter *k* = 1. Importantly, considering the cell cycle, a gamma function (with a large *k*) should make more sense biologically, and using an exponential (or geometric) function might create quite a different pattern of ITH from those simulated with a gamma function (see below). In summary, tumopp is flexible enough to simulate a tumor under various conditions. It not only allows simulations under near identical settings as most previous simulation studies but also exploration of the robustness of any findings by comparison of simulation results with various settings.

## Results

As shown in Table 1, tumopp is much more flexible compared with the four previous models, which arbitrarily explored only limited conditions. Our simulator has a number of options listed in Table 1, which cover almost all settings used in the previous studies. Here, we demonstrate how the different options in tumopp affect the final outcome. In the current work, we used a 3D regular lattice and Moore’s definition of neighborhood to be comparable with previous studies. Essentially identical results can be obtained in a 3D hexagonal lattice, whereas some unrealistic outcomes may be obtained if the von Neumann neighborhood is assumed (see Discussion). First, we give an overview of the results under neutrality (assuming no driver mutations), followed by a discussion of the results with driver mutations.

### Tumor growth patterns and cell genealogy under neutrality

Because the cell division rate should be much larger than the death and migration rates in a tumor, we first ignored the latter two rates. Push method 2 was used because the effect of push methods is negligible on the pattern of tumor growth (but quite large on ITH, as shown in the next section). We first assume that all cells are CSCs (i.e., *p*_*s*_= 1) having the same cell division rate regardless of local density (i.e., constant-rate model). Under this condition, the major factor used to determine the growth curve of a tumor is the shape parameter of the gamma distribution, *k*. We performed simulations with various values of *k*, and typical patterns are shown in Fig. 5. Each simulation run was terminated when the total number of cells reached *N* = 2^14^≈ 16, 000. When *k* = ∞ and all cells undergo cell division at the same time, the tumor grows stepwise (right panel, Fig. 5), and the number of cell divisions experienced (denoted by *v*) is identical for all cells in the final tumor, resulting in a symmetric genealogy with *v* = 14 for all cells (top left genealogy, Fig. 5). As *k* decreases, the variance in *w*_*b, i*_ increases along with the variance of *v*. The other extreme case is *k* = 1 where cell division occurs regardless of the cell cycle, which is the assumption used in most previous studies [9, 17–19]. The growth curve is near exponential, and we observe a substantial variation of *v* in the final tumor (bottom genealogy, Fig. 5). This means that some cells may undergo a large number of cell divisions and some may not.

**Table 1.**
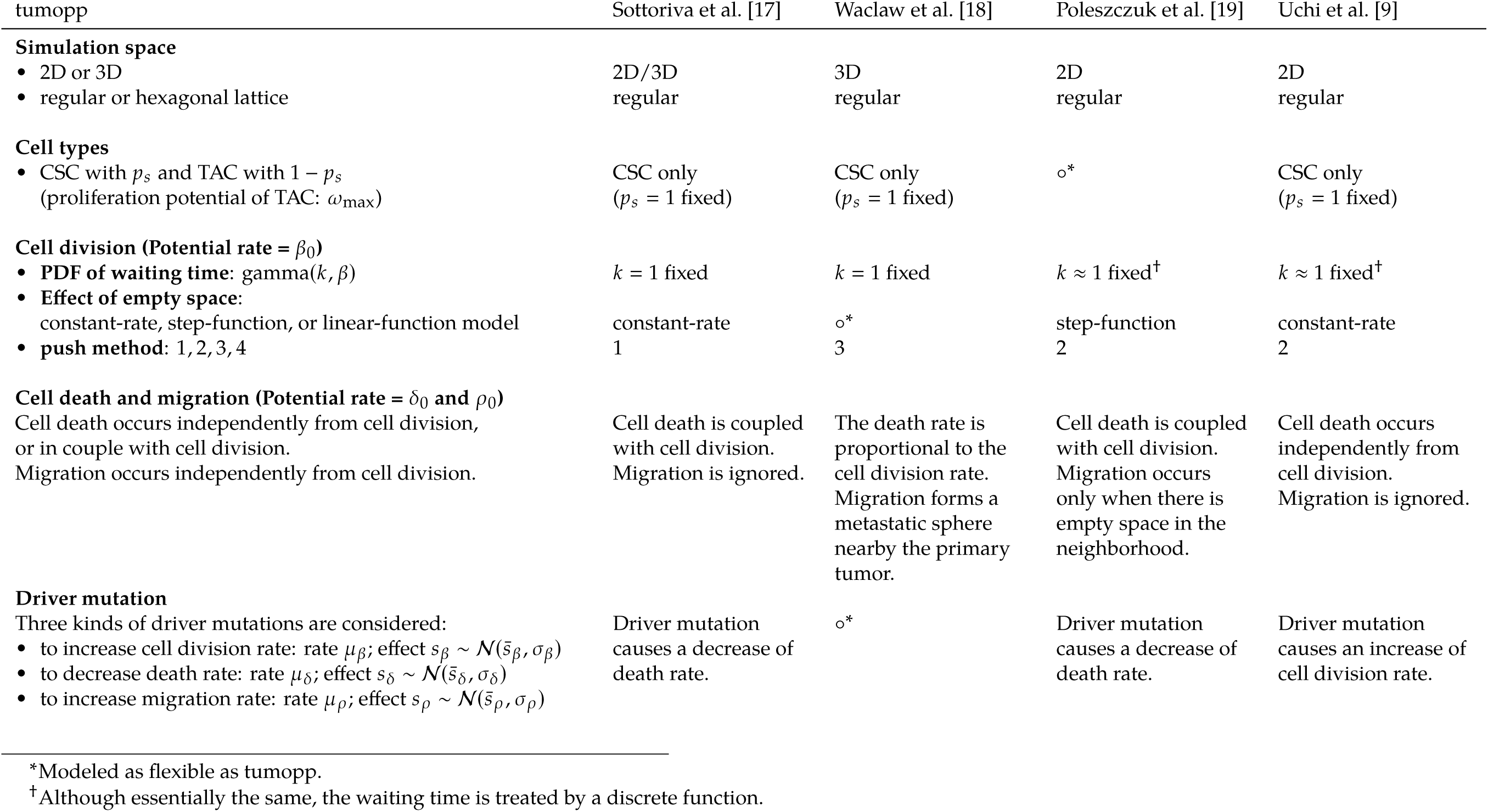
Summary of tummop compared with four previous studies.

**Fig 5.**
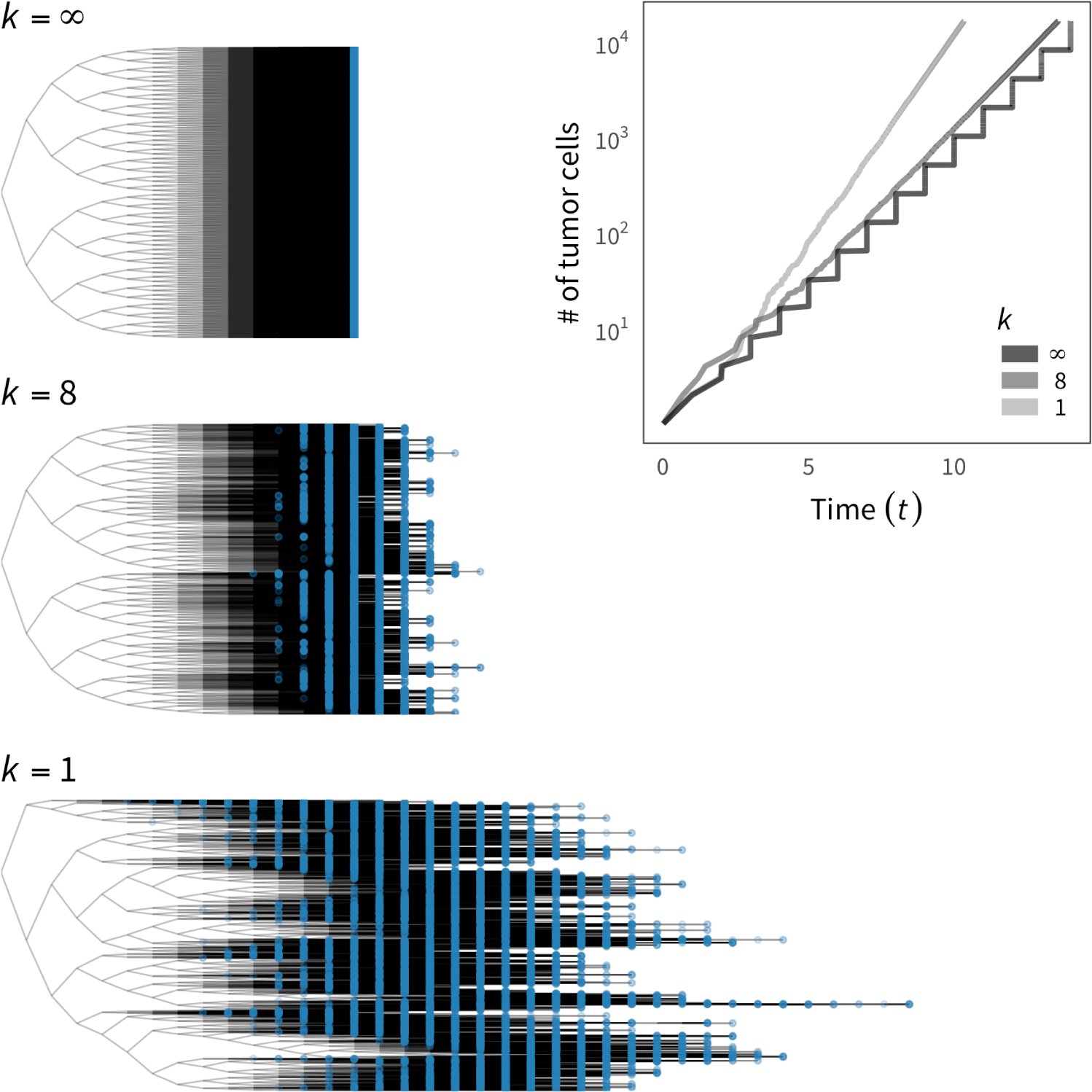
Effect of the shape parameter of the gamma distribution (*k*) on the tumor growth curve and cell genealogy. Three values of *k* = {1, 8, ∞} are used. The cells from the final tumor are represented by blue circles on the genealogies. The constant-rate model is assumed to demonstrate the point.

It should be noted that the growth rate in the right panel of Fig. 5 is negatively correlated with *k*, even when we set an identical birth rate, like β = 1 and *w*_*b*_ = 1 for all cells at birth (or cell division). The growth rate is smallest when *k* = ∞, where the growth curve is deterministically given by *N*_*t*_ = 2^*t*^ because ∆*t* = 1 at any cell division event. When *k* is finite, the growth curve is not deterministic because it involves a random process; the system proceeds by choosing the smallest waiting time, which presumes E(∆*t*) < 1. The growth rate is largest when *k* = 1, where the expected number of tumor cells at time *t* is given by *N*_*t*_=*e*^*t*^.

Fig. 6A shows typical growth curves and genealogies under the constant-rate (blue), step-function (yellow), and linear-function (red) models for E2 that determines how local density affects the cell division rate. The constant-rate model assumes a fixed cell division rate, while the latter two assume the rate as a function of local density. *k* = ∞ is fixed to demonstrate the point because essentially identical results were obtained for other values. The tree on the top with blue nodes for the constant-rate model is the same as the top genealogy in Fig. 5. This figure shows that if the step- and linear-function models are used, competition with neighboring cells is incorporated such that the cell division rate decreases (E2, Eq. 8). This causes a substantial variance in the number of cell divisions per cell (*v*). Consequently, growth under these models (yellow and red, inner panel, Fig. 6A) is slower than that under the constant-rate model (blue, inner panel, Fig. 6A).

**Fig 6.**
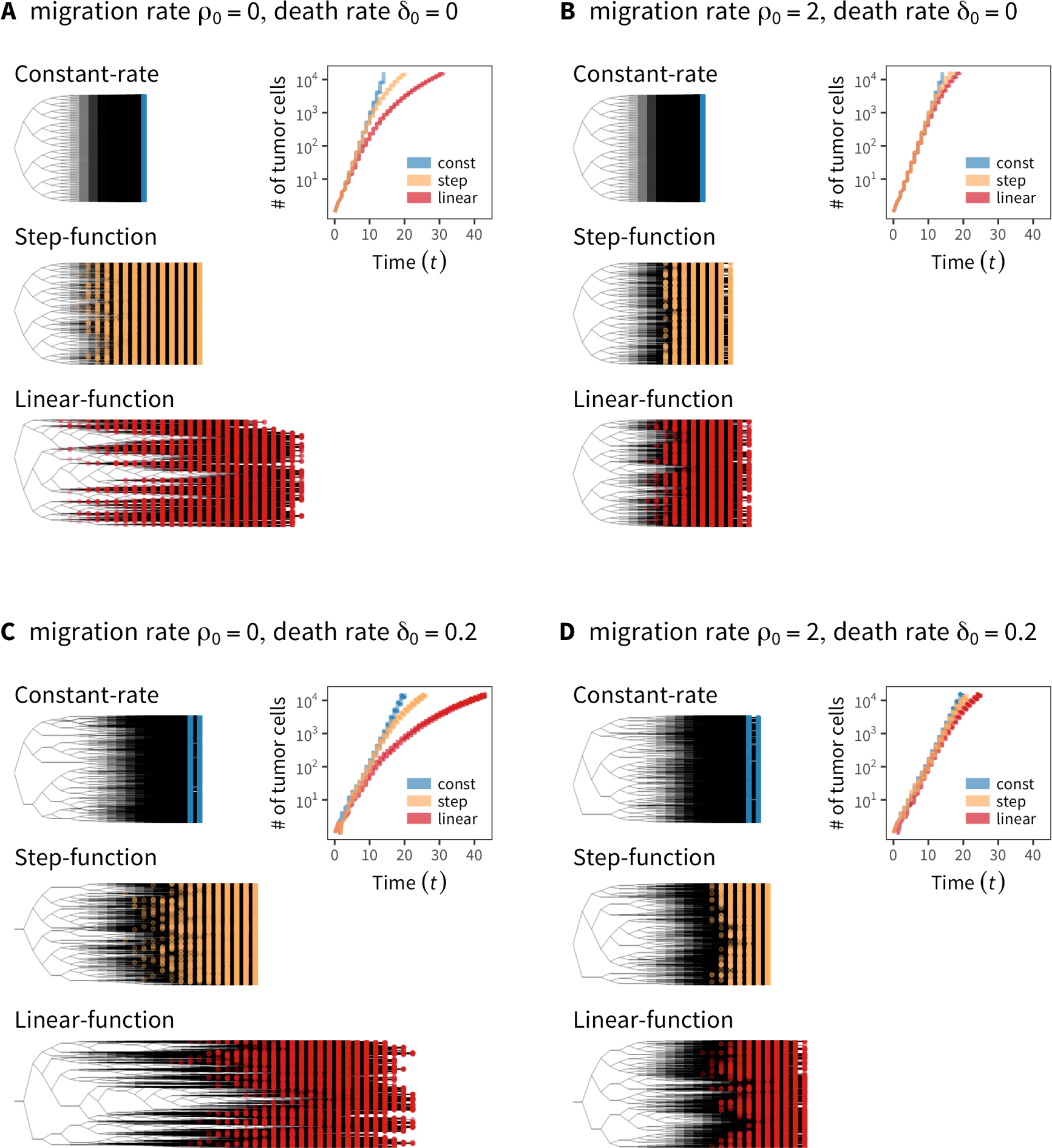
The effect of local density on the tumor growth curve and cell genealogy under the constant-rate, step-function, and linear-function models for E2. Simulation results with (A) no cell death or migration, (B) migration (ρ_0_=0) but no death, (C) death (δ_0_ = 0.2) but no migration, (D) both migration and death (δ_0_=0.2 and ρ_0_ = 2).

This slowing of growth is somewhat diminished when we introduce migration (Fig. 6B). Migration could transfer cells to less crowded sites, thereby resulting in an increase in growth rate (Fig. 6B). This applies to the step- and linear-function models, while the result for the constant-rate model is identical to that in Fig. 6A because it assumes a constant cell division rate regardless of local density. If cell death is incorporated (Fig. 6C), we observe an obvious reduction in growth rate in all three models for E2. Fig. 6D shows the joint work of migration and cell death.

Next, we considered the effect of cell differentiation by additional simulations with the same parameter sets as Fig. 6, except that the assumption of all CSCs is relaxed. Fig. 7 shows the result for the step-function model because we obtained essentially the same result for the linear-function model (the constant-rate model was not relevant here because it allows cell division regardless of the availability of space in the neighborhood). The case wherein no CSCs migrated or died (yellow curve, Fig. 7) is shown as a standard for comparison, which is identical to Fig. 6A; the growth curve with *p*_*s*_ = 0:2 (purple line) illustrates that a CSC undergoes an asymmetric cell division and produces a TAC at probability 1 − *p*_*s*_ = 0.8, and a TAC eventually becomes a TDC after *ω*_max_ = 5 cell divisions. This figure also shows the tumor stopped growing at *t* = 25 because it was completely surrounded by immortal TDCs, thereby creating a barrier that prohibits inside cells from undergoing further divisions. The inner panel of Fig. 7 illustrates this type of situation, where a 2D hexagonal lattice is assumed to demonstrate the point. The dark purple cells with *ω* = 0 are TDCs that completely surround the entire tumor, prohibiting further division of inner cells. This applies only when there is no migration or death so that the barrier will work “forever” once established. If migration or cell death is introduced, the barrier is not permanent or may not even be established (dark and light green lines, Fig. 7). This phenomenon was pointed out in a previous study [19] and is well confirmed in our simulation.

**Fig 7.**
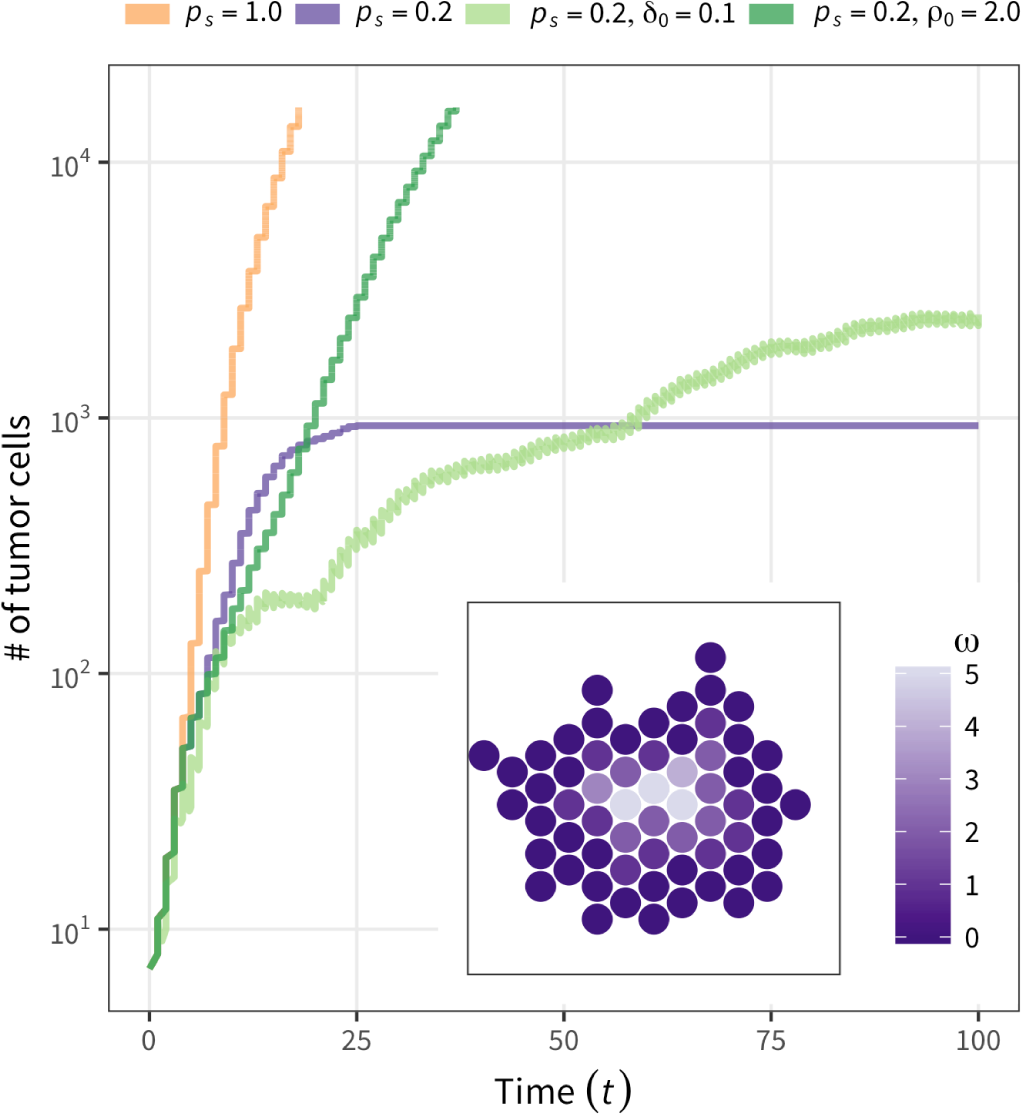
Typical tumor growth behavior when the assumption of all CSCs is relaxed. *p*_*s*_ = 0.2 and *ω*_max_ = 5 are assumed, except for the case of involving all CSCs (*p*_*s*_= 1) for comparison (yellow line). With no cell death or migration (purple line), growth likely stops when the tumor is surrounded by immortal TDCs (inner panel). This effect can be moderated by cell death and/or migration (light and dark green lines).

### ITH and tumor shape under neutrality

The choice of setting in our simulator markedly affects the ITH pattern and shape of the final tumor. Again, we first assumed that no migration or cell death occurs and that all cells are CSCs (*p*_*s*_ = 1 fixed). After performing a large number of simulations under various settings, Fig. 8 shows the observed patterns in eight pairs of E2 models and push methods: 4 push methods under the constant-rate model; 2 push methods under each step- and linear-function model (the behaviors of push methods 2–4 under the step- and linear-function models are identical). For each pair, Fig. 8 presents the results of three independent replicates for two values of *k* (*k* = 1 and ∞). All simulation runs started with a single-cell, and division was allowed until the number of cells hit 10,000; descendants of the first four cells are shown in blue, green, yellow, and red in 3D space (Fig. 8).

One major difference is seen between *k* = 1 and ∞ (left and right halves, Fig. 8): all cells undergo cell division simultaneously when *k* = ∞ (Fig. 5), so the proportion of cell colors is always 25%:25%:25%:25%, and the proportion deviates from this ratio as *k* decreases. This effect is theoretically true, although not visually obvious in Fig. 8. Another difference is how the four colors of cells distribute in 3D space. In the top four rows of the constant-rate model, the four colors of cells are generally intermixed, particularly when push method 1 is employed. This is because cell divisions occur independently of local density in the constant-rate model, and new cells are placed by randomly pushing other cells toward the surface. In contrast, in the step- and linear-function model rows, cells of the same color are more likely located close to one another, making clusters of cells with the same color. This is particularly notable with push methods 2–4, in which a new daughter cell is always placed at an adjacent site if space is available so that closely related cells tend to be located close together.

This pattern is better documented by looking at the relationship between *F*_*ST*_ and physical distance. *F*_*ST*_ is a measure of relative population differentiation at the DNA level. We computed *F*_*ST*_ for a number of pairs of random subregions with size 20 cells from a single tumor. Note that *F*_*ST*_ was computed based on the branch lengths on the genealogy rather than making genetic data by distributing passenger (neutral) genetic markers (e.g., single nucleotide polymorphisms) across the genome; therefore, this *F*_*ST*_ is the expected value when there are an infinite number of markers. The physical distance was computed as the Euclidean distance between the central cells of two subregions. Fig. 9 shows the relationship between *F*_*ST*_ and physical distance for all simulated tumors in Fig. 8. As expected, *F*_*ST*_ and physical distance are more positively correlated when the step- and linear-function models are used.

The shape (morphology) of the final tumor also varies depending on the models for E2 and push methods. Tumors in most cases are more like spheres. Exceptions include cases with push methods 3 and 4 under the constant-rate model, where the final tumors are angular with quite flat surfaces. In these specific cases, there could be a systematic pressure to keep flat surfaces because hollows are quickly flattened by filling new cells from the inside. Other than these exceptional cases, there is some quantitative variation in the deviation from a sphere. It should be noted that irregular morphologies with dramatic deviation from a sphere may correlate with tumor invasiveness [20, 21, 31–33]. It seems the tumor shape is most distorted in the linear-function model. This is because the linear-function model assumes high rates of division for cells with many empty sites in the neighborhood, which largely applies to cells that form outshoots on the surface. As a consequence, such an outshoot likely grows to be a lump, thereby resulting in a marked deviation from a spherical shape. This also explains the observation that *F*_*ST*_ and physical distance are most strongly correlated in the linear-function model.

**Fig 8.**
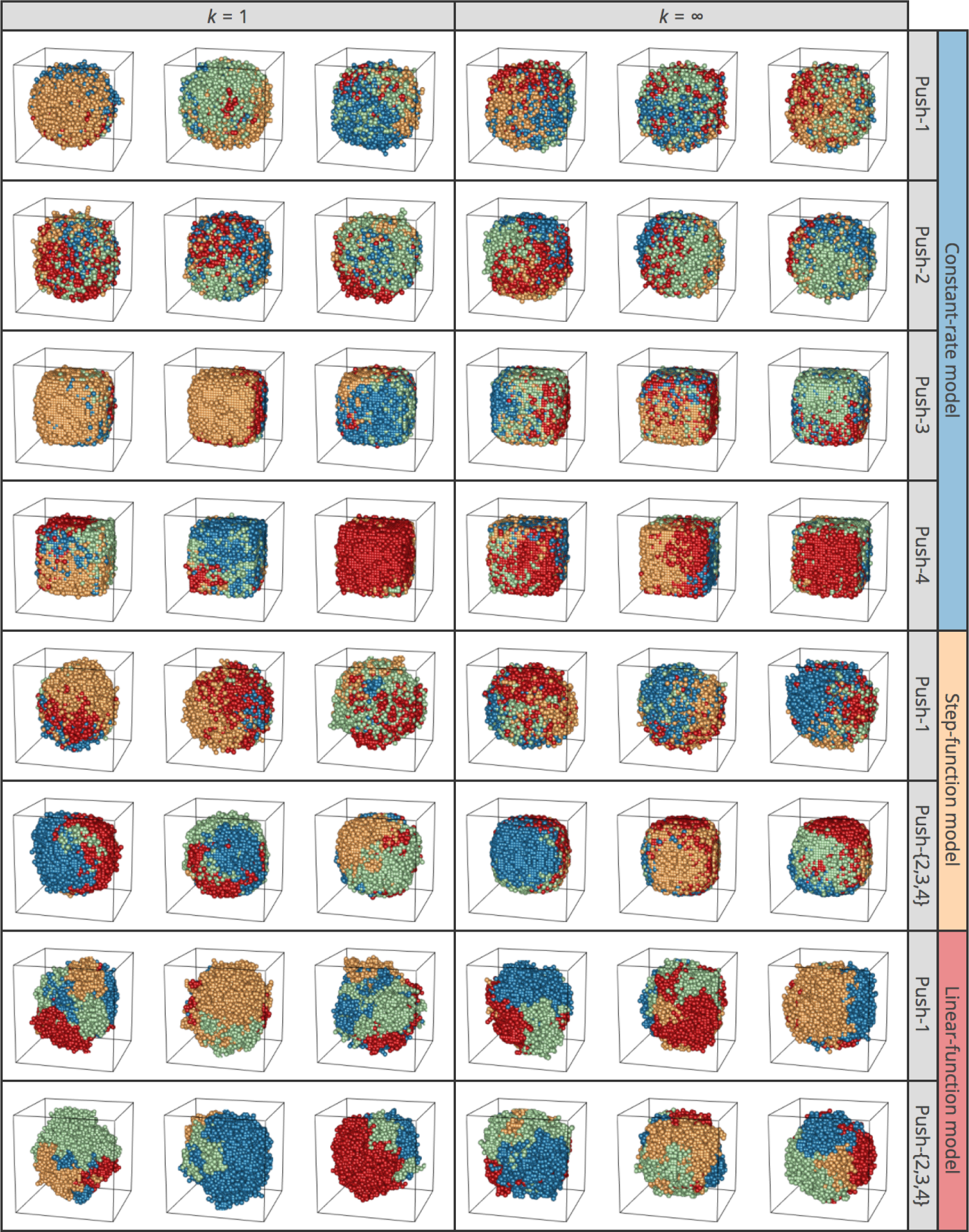
3D structures of simulated tumors for push methods 1–4 under constant-rate, step-function, and linear-function models under neutrality. Results for *k* = 1 and ∞ are shown. Descendants from the first four cells in each simulation run are shown in blue, green, yellow, and red. The results of three independent runs are shown for each setting.

**Fig 9.**
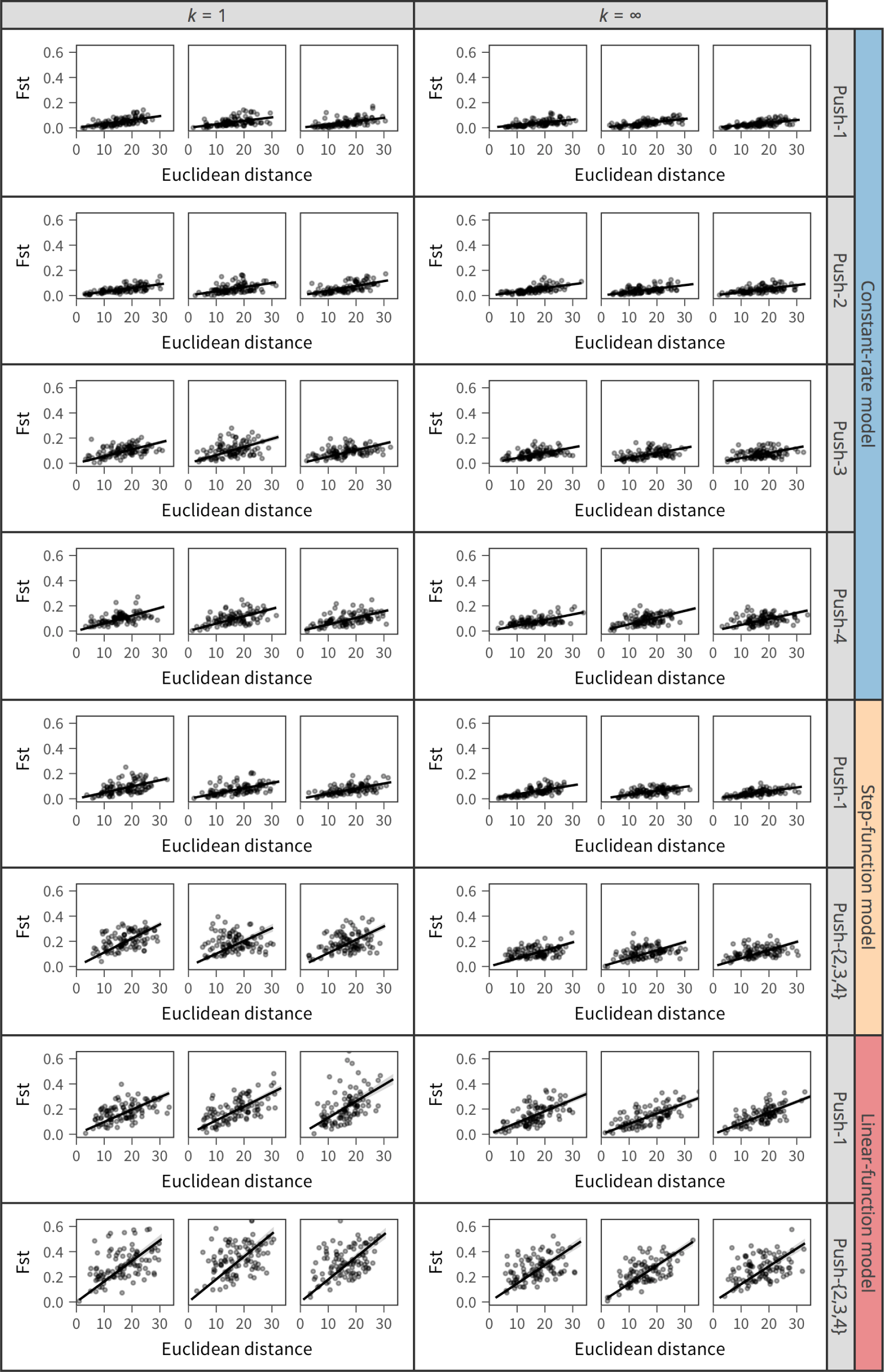
Correlation between *F*_*ST*_ and physical distance measured by Euclidean distance. The simulated tumors shown in Fig. 8 are used.

Fig. 10 explores the effect of cell death and migration. We show only results for *k* = ∞ because essentially the same results were obtained for other values of *k*, including *k* = 1. The plots in the left quarter were obtained with the same parameter sets as those in the right half of Fig. 8. It appears that the effect of adding cell death alone (ρ = 0.2) may be small, while migration tends to create more distorted tumors, with more intermixing of the four cell colors (right half, Fig. 10). It is also notable that we observe a number of outshoots on the surface when migration is included.

**Fig 10.**
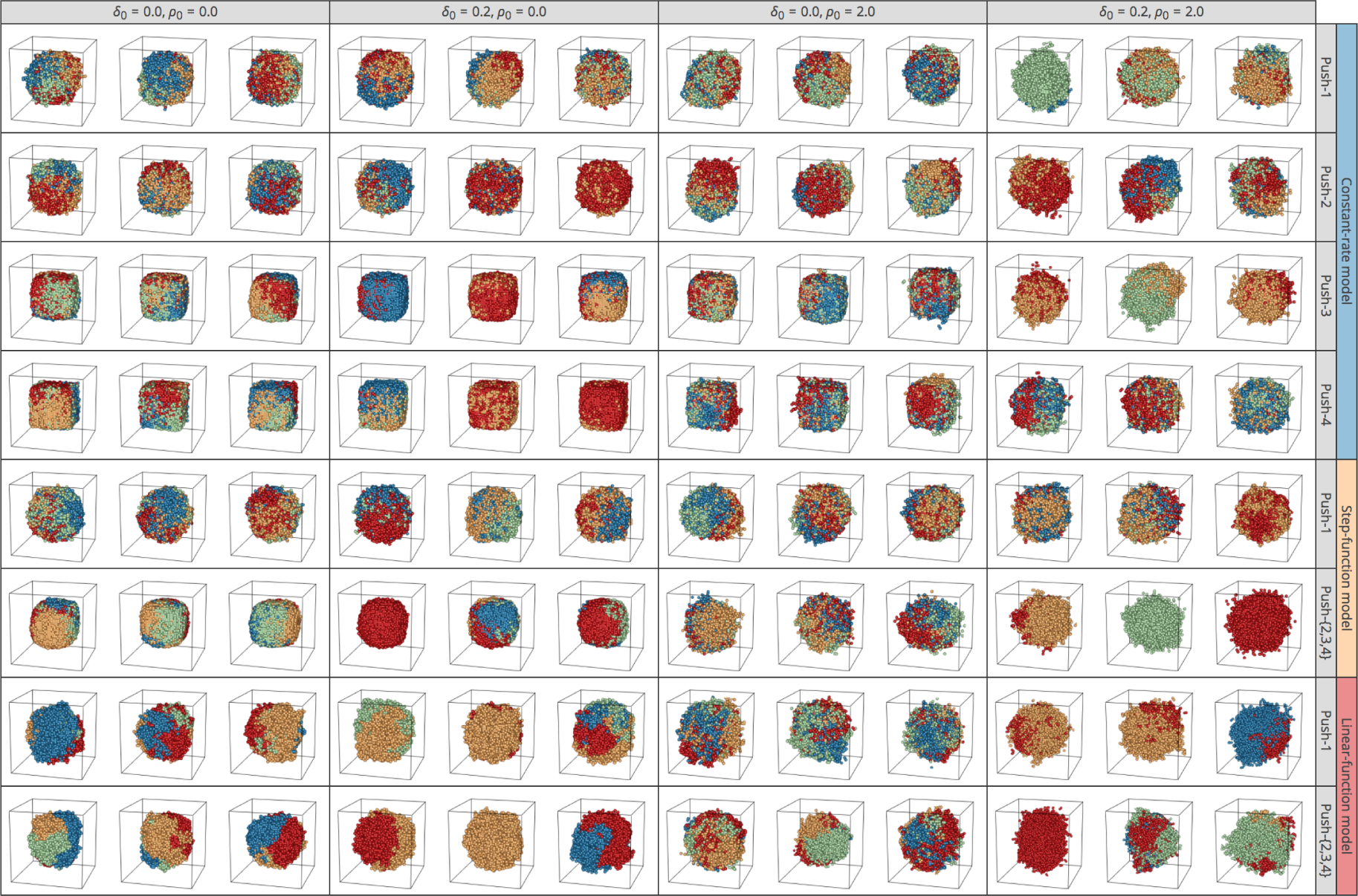
Effect of cell death and migration on the 3D structures of simulated tumors for push methods 1–4 under constant-rate, step-function, and linear-function models. *k* = ∞ is assumed. Descendants of the first four cells in each simulation run are presented in blue, green, yellow, and red. The results of three independent runs are shown for each setting.

In Fig. 11, we further relaxed the assumption of all CSCs. We used two values for the cell differentiation parameter *p*_*s*_ = 0.6, 0.2), withω_max_= 5 and 10. We show the results when the step-function model and push method 2 are assumed because essentially the same results were obtained for other settings. The top row of Fig. 11 shows the result for the case involving all CSCs, which was obtained by simulations with the same parameter sets as the sixth row of Fig. 10. The most marked effect of *p*_*s*_ is that tumor growth could stop when it was surrounded by TDCs, as demonstrated in Fig. 7. This effect is well observed particularly when *p*_*s*_ is small (i.e., *p*_*s*_= 0.2),ω_max_ is large, and migration is not allowed (ρ = 0.0) (see Poleszczuk et al. [19]).

**Fig 11.**
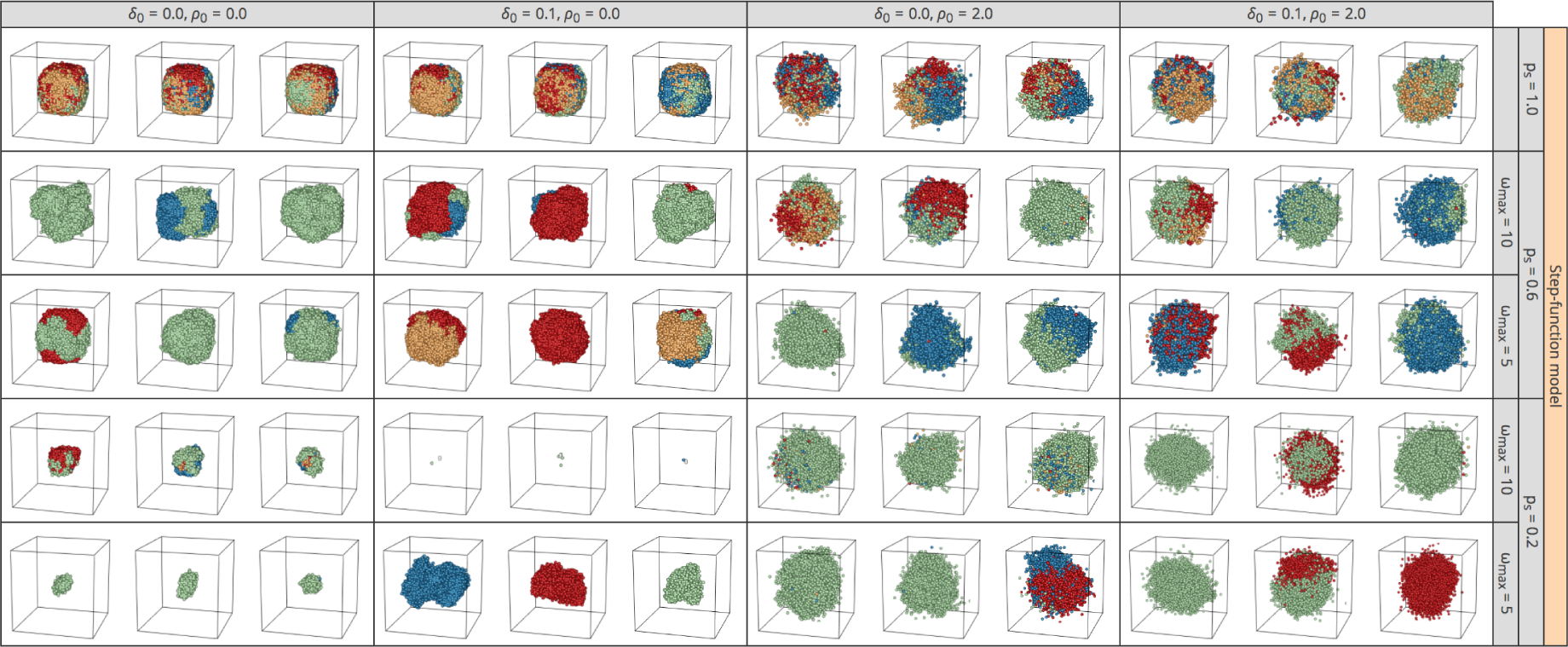
Effect of nonstem cells on the 3D structures of simulated tumors. Results for push method 2 under the step-function model are shown. *k* = ∞ is assumed. Descendants from the first four cells in each simulation run are presented in blue, green, yellow, and red. The results of three independent runs are shown for each setting.

### Effect of driver mutations

Three kinds of driver mutations are implemented in tumopp, those that increase the cell division rate, decrease the cell death rate, and increase the migration rate. Here, we focused on the first type of driver mutations that increase the cell division rate because the effects of the other two kinds of driver mutations are relatively simple (data not shown). If driver mutations are assumed to decrease the death rate, the major effect is slowed tumor growth, and driver mutations that increase the migration rate would create a more intermixed spatial distribution of cells of different genotypes.

There would be two extreme cases for driver mutations that increase the cell division rate: (i) driver mutations with small effects arising frequently (Fig. 12) and (ii) a driver mutation with a large effect occurs only once (Fig. 13). We show some simulation results for these two cases with relatively simple settings to demonstrate this point. Cell death and migration are ignored (δ_0_ = 0, ρ_0_ = 0), and all cells are CSCs (*p*_*s*_ = 1), which is the same setting used in Fig. 6A, with a slight modification: *k* = 100 is assumed instead of *k* = ∞. This modification was made because *k* = ∞ predicts all cells undergo cell division simultaneously and that the cell number grows stepwise (ladder line, Fig. 5), which is not suitable if we want to introduce a driver mutation at an arbitrary time point specified by the size of tumor (*N*_µ_). This applies to the simulation for (ii).

**Fig 12.**
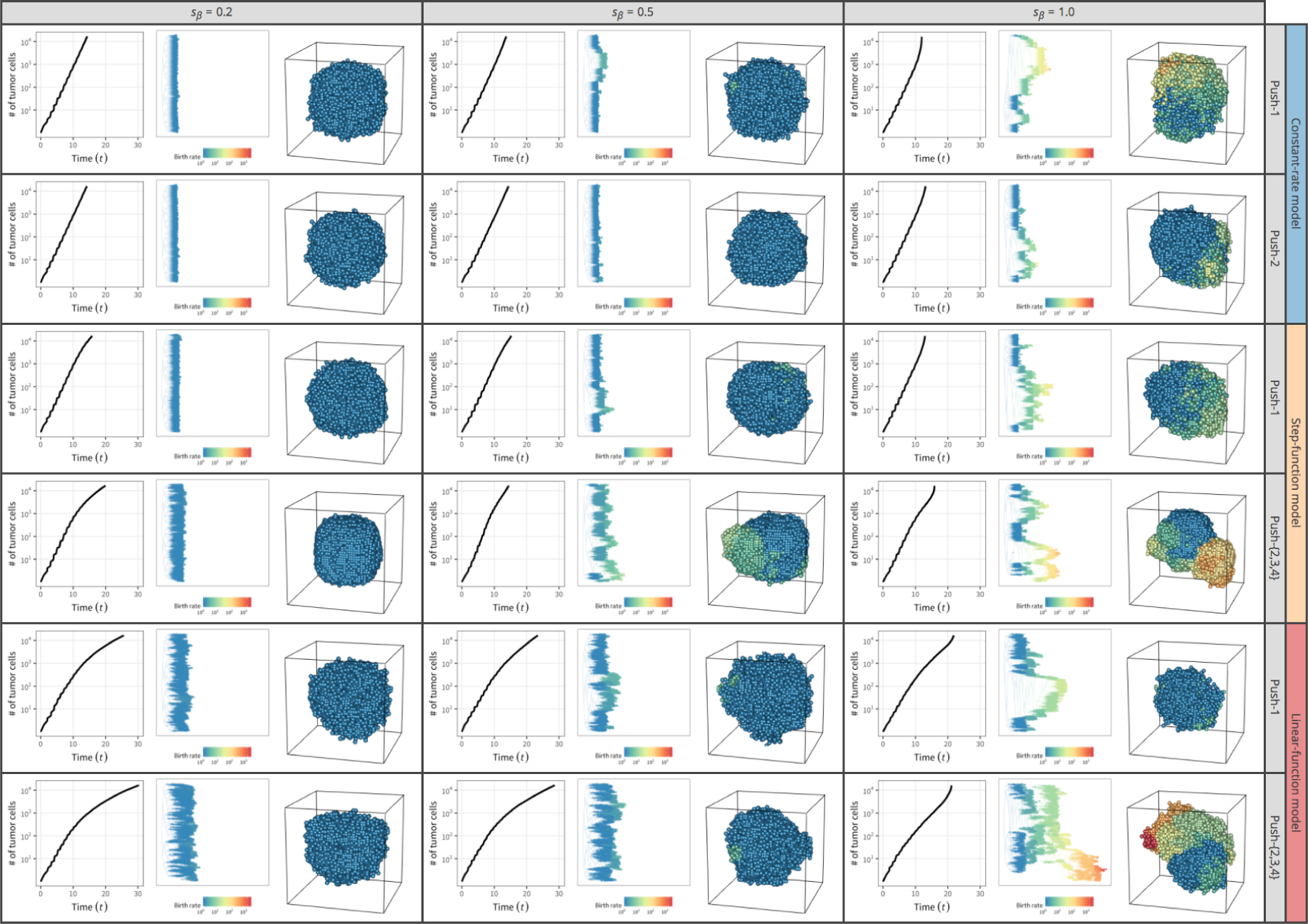
3D structures of simulated tumors with frequent weak driver mutations. Results for push methods 1 and 2 under constant-rate, step-function, and linear-function models are shown; *k* = 100 is assumed. The colors of cells represent their cell division rates, scaled from blue to red. The results for one simulation run are shown for each setting.

The effect is quite different between the cases (i) [Fig. 12] and (ii) [Fig. 13]. In the simulation for case (i), weak driver mutations were assumed to occur quite frequently with parameters µ_β_;0:005, 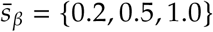
, and ρ_β_ =0. Fig. 12 shows the results for push methods 1 and 2 under the constant-rate, step-function, and linear-function models. The results for push methods 3 and 4 with the constant-rate model are not shown because they are quite similar to those of push method 2 (push methods 2–4 assume the same behavior under the step- and linear-function models). In Fig. 12, cells are shown such that the cell division rate is scaled in color, from blue (β = 1, default rate) to red. Under all settings, it is clearly demonstrated that as average intensity of driver mutations (*s*¯β) increases, the growthrate increases due to the cells that have acquired driver mutations. Cells with driver mutations likely undergo more cell divisions and make a cluster on the surface.

**Fig 13.**
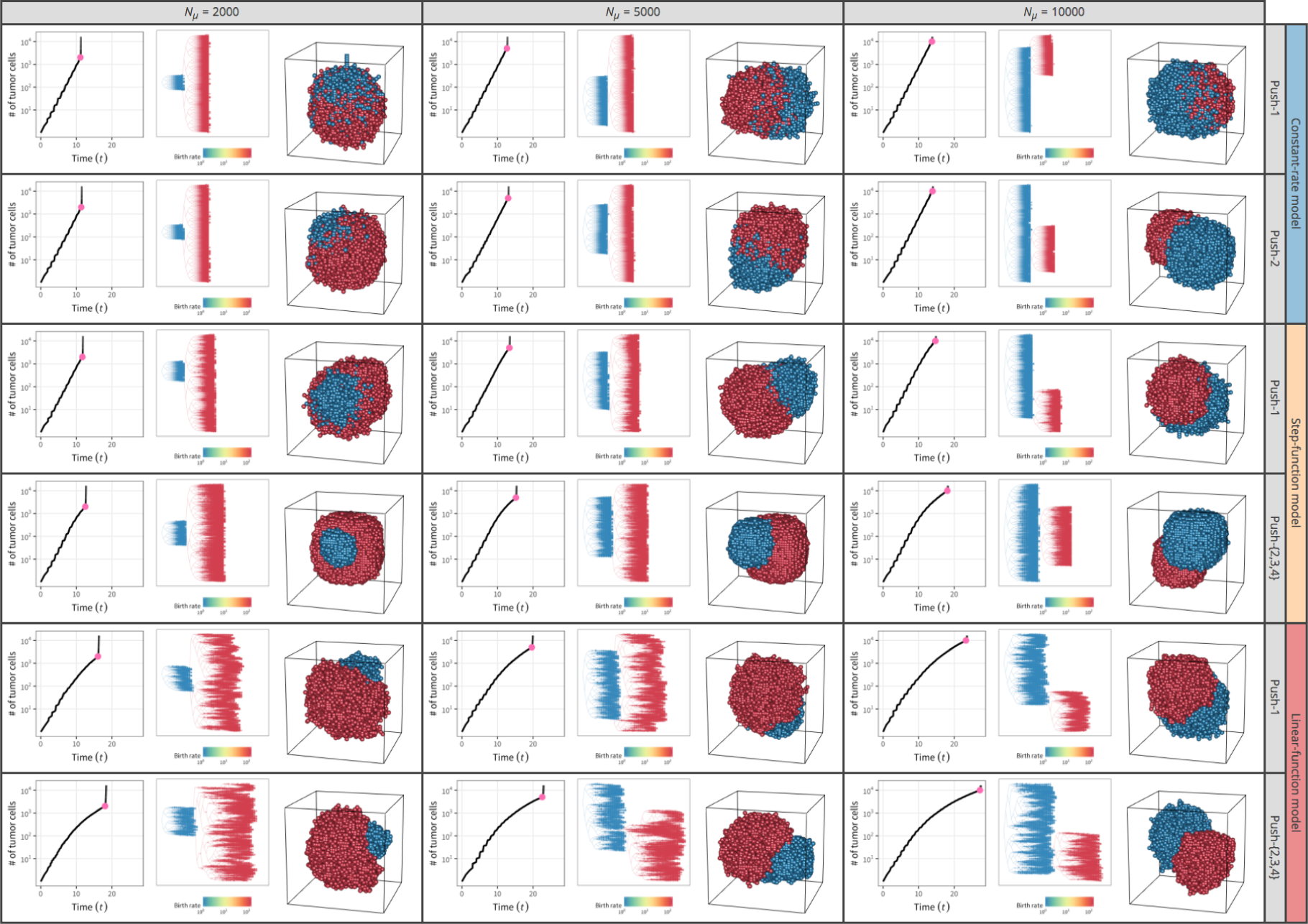
3D structures of simulated tumors with a single strong driver mutation. Results for push methods 1 and 2 under constant-rate, step-function, and linear-function models are shown; *k* = 100 is assumed. The cells with the driver mutation are in red, while the others are in blue. The results for one simulation run are shown for each setting. The time point when the driver mutation was introduced is shown by a pink circle on the growth curve.

With 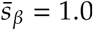
, the cell division rate increases to β > 200 (orange to red), creating quite skewed tumor shapes with accelerated growth rates. Particularly for push methods 2–4 with the step- and linear-function models, the 3D structure of the tumors is complicated because the step- and linear-function models assume the cell division rate is on average higher on the surface.

Fig. 13 considers the other extreme case (ii), where a single, very strong driver mutation is introduced arbitrarily. During each simulation run, rather than setting the driver mutation rate, we arbitrarily introduced a strong driver mutation with *s_β_* = 99 when the number of tumor cells reached *N*_µ_ ={2000, 5000, 10000}. An *s*_β_= 99 means that a single mutation caused an increase in cell division rate 100 times as high as the original value. Fig. 13 shows the results for push methods 1 and 2 with constant-rate, step-function, and linear-function models. Even with very low initial frequencies (i.e., {1/2000; 1/5000; 1/10000} for *N*_µ_ {2000, 5000, 10000}, respectively), the cells with the driver mutation (red, Fig. 13) grow dramatically, resulting in an immediate increase of the total number of cells. It seems that the red cells with the driver mutation likely result in a distinct cluster particularly for push method 2 with the step- and linear-function models, whereas red and blue cells are to some extent intermixed in the constant-rate model.

## Discussion

Herein, we developed a simulator named tumopp that generates ITH patterns. Thus far, ITH simulations have been conducted in several previous studies; however, the model settings used varied (Table 1). This means that only limited conditions were explored in each study. Motivated by this issue, we developed tumopp to be as flexible as possible so that all four previous models could be included and making it extremely useful for exploring the effects of model and parameter settings. Variations in the model settings include how the cell division rate is determined, how daughter cells are placed, and how driver mutations are treated. Moreover, to account for the cell cycle, we introduced a gamma function for the waiting time involved in cell division, while all previous studies adopted simple decreasing (e.g., exponential) functions (Fig. 3). In our model, the shape of the gamma distribution can be specified by parameter *k*, and a *k* = 0 gives an exponential distribution whereas *k* = ∞ assumes that all cells undergo division simultaneously.

Moreover, tumopp uniquely implements a hexagonal lattice, which we believe is biologically more reasonable because the distance to all neighbor cells is identical so that there is only one definition of the neighborhood (Fig. 2). S1 Fig briefly shows simulated tumors in a 3D hexagonal lattice with the same setting as those used in Fig. 8. We suggest using a hexagonal lattice for future work although we here used a regular lattice to be comparable with the previous studies. Although tumopp implements two definitions of the neighborhood in a regular lattice, we used the Moore neighborhood as in previous studies. The von Neumann neighborhood has not been used often and can create diamond-like tumors, which is obviously an unrealistic morphology (S2 Fig).

Using tumopp, we investigated how model and parameter settings affect tumor growth curves and ITH. We found that *k* (shape) for the waiting time mainly specifies the growth curve (Fig. 5). Moreover, the combined effect of local density on the cell division rate (constant-rate, step-function, and linear-function models), the method to place new cells (push methods 1–4), and cell differentiation plays a role in tumor growth (Fig. 6).

Various patterns in the shape of tumor and ITH arose depending on the model setting. The methods used to determine the cell division rate (i.e., constant-rate, step-function, and linear-function models) and those to place new cells (i.e, push methods 1–4) had a major effect. Under the constant-rate model with push method 1, all cells undergo cell division at a constant rate regardless of local density, and new cells are placed randomly pushing out pre-existing neighbor cells. This behavior makes shuffled patterns of ITH with weak isolation by distance (Figs. 8 and 9). By contrast, under the linear-function model with push methods 2–4, the cell division rate is higher when more space (empty sites) is available in the neighborhood, which generally applies to cells near the surface (particularly to cells that constitute outshoots from the surface); new cells are placed to fill the empty space without pushing existing cells. This setting likely creates a biased complex shape of tumor with clusters of genetically closely related cells, resulting in strong isolation by distance.

The effects of driver mutations were implemented by increasing the cell division rate, decreasing the death rate, and increasing the migration rate. Our simulation demonstrated that the effect of driver mutations on ITH would be remarkable when introduced to increase the cell division rate, especially when driver mutations with large effects are involved. Although this mode of driver mutation was implemented in Waclaw et al. [18] and Uchi et al. [9], the effects on ITH and tumor morphology were not fully explored. Tumor growth dynamics with various kinds of driver mutations would be an intriguing subject for future study. It would also be interesting to involve environmental changes, which can be easily incorporated in tumopp. For example, chemical agents would cause a dramatic reduction in the size of the cancer cell population, and a regrowth of remaining resistant cells might occur. Simulations with such environmental changes would give insights into the behavior of tumors after medical treatments.

Although tumopp may take a considerable amount of time to simulate very large tumors, this problem may be solved to some extent if the tumor is assumed to consist of compartments; for example, glands in a colorectal tumor, as pointed out by Sottoriva et al. [17, 34]. Glands proliferate through gland fission [35], and each gland is almost a clonal population of the cells originated from a few CSCs [36–38]. If so, when simulating a tumor, a compartment can be treated as a single unit (cell). A compartment-based simulation would involve much less computational load than a cell-based simulation.

Our work demonstrates that extremely variable patterns of ITH can be created even under neutrality, depending on the model setting. This suggests a caveat in analyzing ITH data with simulations with limited settings because another setting might predict a different ITH pattern, which could result in a different conclusion. For example, Sottoriva et al. [17] investigated ITH in colorectal tumors by sequencing a number of glands from single tumors. They found that cancer cells with similar genetic backgrounds were observed on both the left and right sides of the tumors. This observation led the authors to conclude that mutations that arose in early stages spread during growth, and they confirmed that such intermixed tumors can be generated by simple simulations assuming push method 1 with the constant-rate model in our framework. Our simulations agree that this setting produces intermixed tumors but not with other settings, such as push methods 2–4 with the linear-function model. Thus, we suggest that simulation setting be carefully chosen, and deep understanding of the typical behavior of cancer cells is important. Otherwise, it is important to carry out simulations under various conditions to confirm or verify the results. For this purpose, tumopp will be very useful, and the source code is available on GitHub (https://github.com/heavywatal/tumopp).

## Supporting Information

**S1 Fig.**
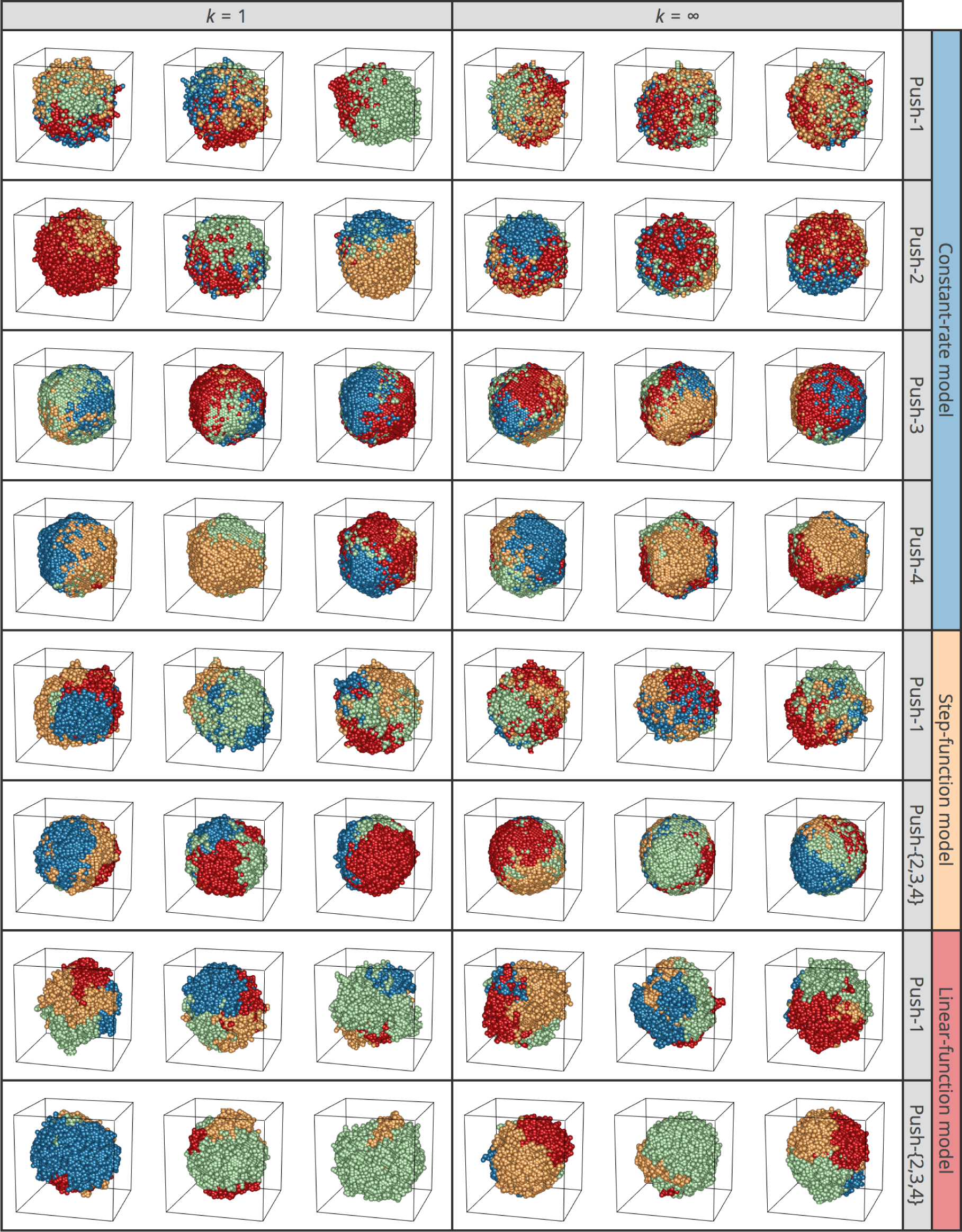
3D structures of simulated tumors in a hexagonal lattice. All parameters except for the lattice/neighborhood are the same as those in Fig. 8.

**S2 Fig.**
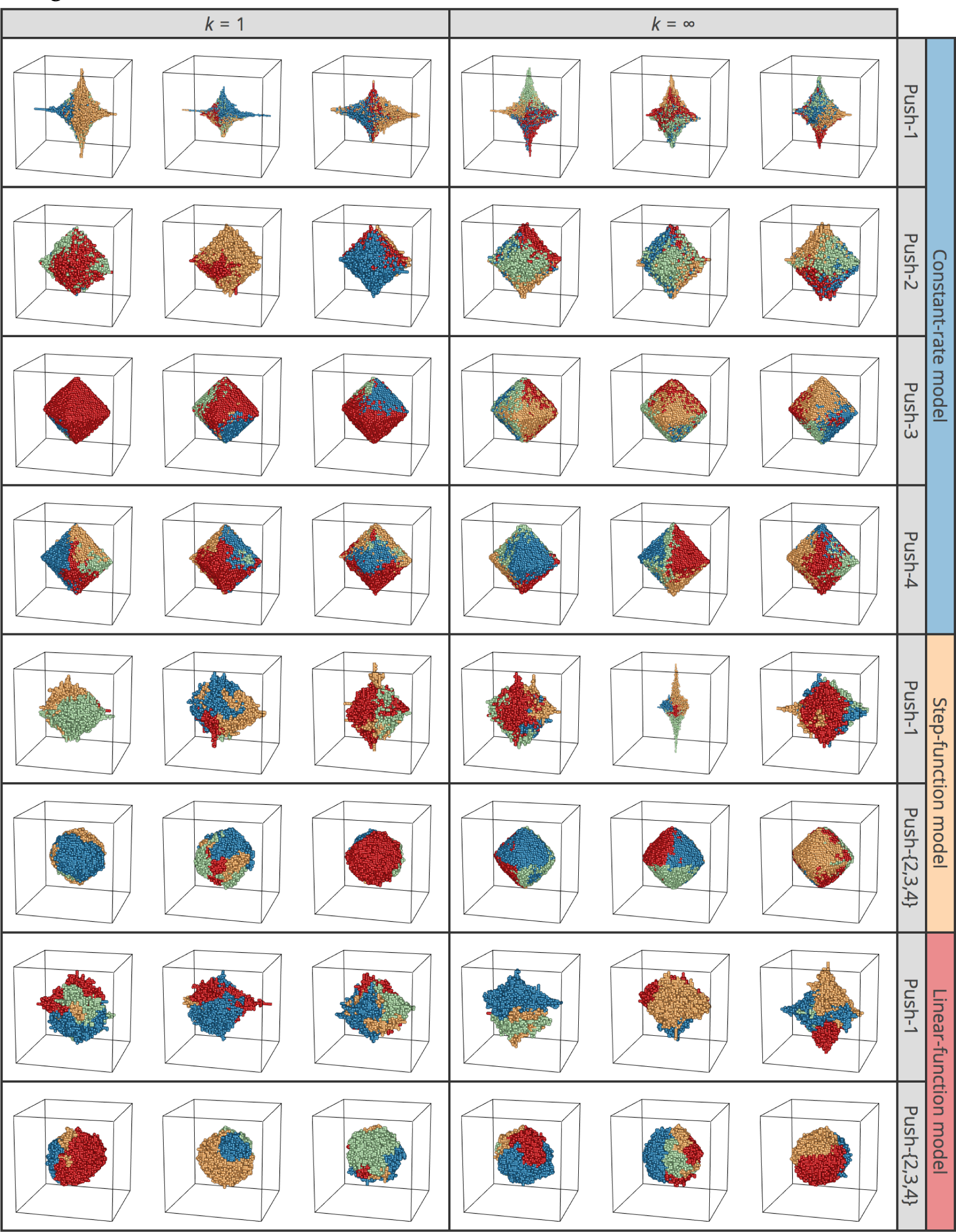
3D structures of simulated tumors assuming the von Neumann neighborhood in a regular lattice. All parameters except for the lattice/neighborhood are the same as in Fig. 8.

## Acknowledgments

We thank Ryuichi P. Sugino and Kazuki Takahashi for valuable discussion on cancer evolution.

